# Apparent digestibility and nutritional performance of bakery waste produced in Punjab Pakistan for *Labeo rohita* fingerlings

**DOI:** 10.1101/2024.10.29.620837

**Authors:** Syed Shafat Hussain, Noor Khan, Fayyaz Rasool, Simon John Davies, Saima Naveed

## Abstract

This study aimed to examine the impact of bakery waste supplementation on the nutrient digestibility and performance metrics of rohu (*Labeo rohita*) juveniles. Fish were fed with seven different diets containing bakery waste products (biscuits, cream cake, bread, and dry rusk) in combination with rice polish and wheat flour. Chromic oxide (1%) was added to the diet as an inert marker to measure nutrient digestibility. The results indicated a significant difference (p < 0.05) in the digestibility of crude protein, lipid, gross energy, and dry matter among the dietary groups. Fish fed with rice polish had the lowest apparent digestibility value of protein (16.13 ± 1.86), whereas fish fed wheat flour had the highest (77.70 ± 0.05). Moreover, it was shown that fish fed rusk waste had better growth. It has been concluded that bakery wastes could be used as a cheaper fish feed ingredient source for the manufacturing of cost-effective feed for *L. rohita*. This investigation was discussed within the context of providing cost effective ingredients for sustainable aquaculture production in Pakistan.

## 1. Introduction

*Labeo rohita* is the most widely cultivated freshwater fish species in Pakistan, it requires accurate data on the nutrient digestibility coefficients of new alternative ingredients. This information will help to sustain its cultivation and meet the protein requirements for economic feasibility and accurate feed formulation [1]. As the most preferred fish in Punjab, Pakistan, *L. rohita* is predominantly cultured intensively in monoculture as well as in polyculture systems. These characteristics make it an ideal candidate for evaluating its specific feed requirements and efficiency with a variety of ingredients, including bakery wastes [2]. As the aquaculture industry is expanding very rapidly, the demand for grains increases, leading to higher prices for both human and animal use. Supplementing fish feed with plant sources is an effective way to provide essential nutrients and reduce reliance on fish meal, thereby partially meet the nutritional needs of fish [2].

It is essential to investigate the digestion and utilization values of feedstuffs, rather than merely analyzing their protein content through chemical analysis. Assessing the actual efficiency of feedstuffs in relation to fish growth requires determining their digestibility [2]. Understanding the digestibility of nutrients in feedstuffs not only maximizes the growth of farmed fish but also reduces costs and minimizes feed waste excreted by the fish [3]. The most pivotal factor evaluating the viability of feedstuff is digestibility, which is the part of ingredients, supplements or energy in the consumed feed that is not removed in the excrement [4]. In order to ascertain the possible nutritional value of substitute feed components and more specialized diets for various fish species, feedstuff digestibility testing is a necessary first step. [5].

ADCs (apparent digestibility coefficients) for protein, fat, gross energy, and dry matter of animal protein sources are indirectly measured by using Chromic oxide (Cr_2_O_3_) [6,7] is a standard technique in animal and fish nutrition studies. Although carbohydrates are inexpensive sources of energy for animals, various fish species digest them at varying rates and assimilation. Carps such as Rohu (*L. rohita*) can use up to 43 percent of their diet’s without experiencing negative health effects [8]. Bakery waste is one of the most economical and alternative resources that are required to solve the issue. Urban regions are hubs of bakery waste where they are generated in large quantities. Because they are composed of grains, bakery wastes or byproducts are nutrient-dense and have a similar composition to grains.

Therefore, the prime purpose of our experiment was to examine the effects of feeding various types of bakery waste on their nutritional digestibility and growth performance of rohu (*L. rohita*) fingerlings. The knowledge gathered from this study could be very helpful in formulating fish feed that is both economical and environmentally beneficial.

## 2. Materials and Methods

### 2.1. Experimental fish and study site

Four hundred and twenty *L. rohita* juveniles in triplicate were randomly assigned to rectangular tanks (435 L capacity) with a stocking density of twenty fish each, after being procured from the Department of Fisheries and Aquaculture, UVAS, Pattoki. The fish were given two weeks to become used to the new experiment conditions before the 90-day study officially began. During the trial, the juveniles were kept disease-free and in good health.

### 2.2. Water quality

Every tank had aeration, and the water’s ideal range for *L. rohita* was maintained in terms of temperature, dissolved oxygen content, and TDS.

### 2.3. Experimental diets

The Lahore region’s commercial factories and markets supplied the feed ingredients. To get the desired particle size, these substances were crushed and finely ground. After blending the dry components for ten to twenty minutes with an electric mixer, distilled water was added gradually. In order to measure the digestibility, 1% of Chromic oxide was added to the meals. Seven diets were formulated with one control diet (fish meal reference diet) and the remaining diets were having partial (30%) inclusion of different types of ingredients (bread waste, dry rusk waste, biscuit waste, cream cake waste, rice polish, and wheat flour) each with a fish meal containing 30% of CP. A bench-top extruder was used to extrude the moist dough, and the dry pellets that were produced were kept at -20°C. Over the course of the 12-week trial, fish were fed at a feeding rate of 2% of their live wet weight, with weekly adjustments made based on biomass levels. The proximate analysis of the different bakery ingredients are displayed in table 1, while formulation of different experimental diets are listed in table 2.

**Table 1.**
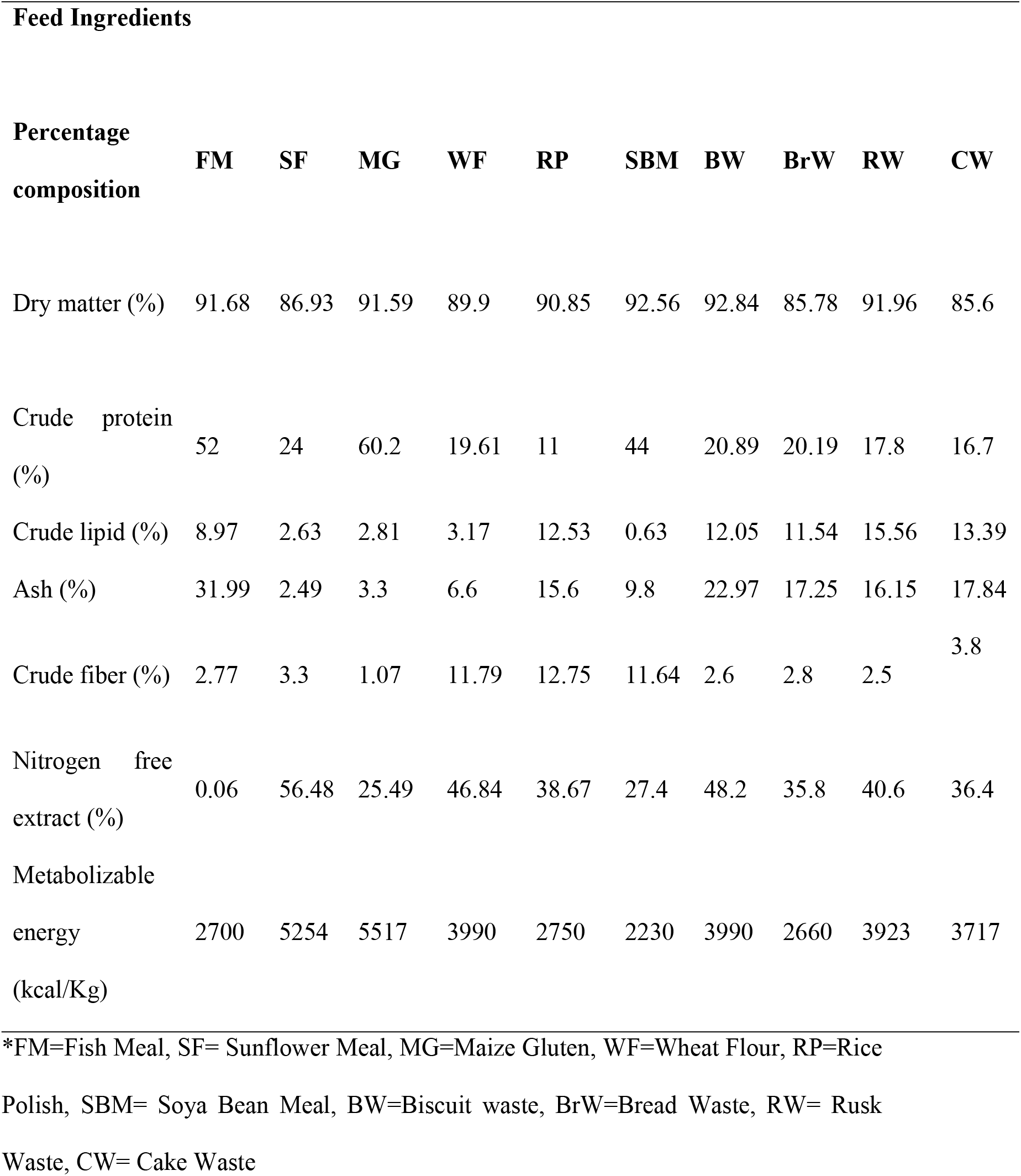
Proximate composition of feed ingredients including different bakery waste.

**Table 2.** Ingredient composition of experimental diets.

| Items | Z | A | B | C | D | E | F |
| --- | --- | --- | --- | --- | --- | --- | --- |
| Fishmeal | 99 | 69 | 69 | 69 | 69 | 69 | 69 |
| Wheat flour | 0 | 30 | 0 | 0 | 0 | 0 | 0 |
| Cake waste | 0 | 0 | 30 | 0 | 0 | 0 | 0 |
| Rice Polish | 0 | 0 | 0 | 30 | 0 | 0 | 0 |
| Biscuit waste | 0 | 0 | 0 | 0 | 30 | 0 | 0 |
| Bread waste | 0 | 0 | 0 | 0 | 0 | 30 | 0 |
| Rusk waste | 0 | 0 | 0 | 0 | 0 | 0 | 30 |
| Chromic oxide | 1 | 1 | 1 | 1 | 1 | 1 | 1 |
| Total | 100 | 100 | 100 | 100 | 100 | 100 | 100 |

\*Z=Control Diet-Fish Meal, A= Wheat Flour, B= Cake Waste (CW), C= Rice Polish, D= Biscuit Waste, E= Bread Waste, F=Rusk Waste.

### 2.4. Fecal collection protocol

After an hour of feeding, the leftover feed pellets and other debris were taken out. From the second week onwards, after two hours of feeding, fecal material was gently collected using siphon mechanism from the tanks onto a bolting silk cloth. After that, it was washed with distilled water and quickly dried with the help of filter paper and stored at -20 °C.

### 2.5. Growth parameters and feed utilization

The apparent digestibility coefficient of dry matter and crude protein of the experimental diets were calculated according to Smith and Tabrett [9] as follows:

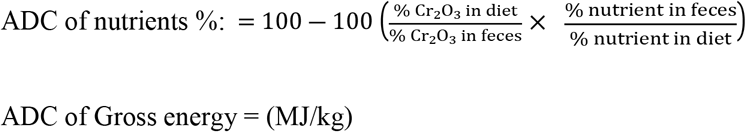

ADC of gross energy was calculated using gross energy data (MJ/kg) instead of % nutrient data.

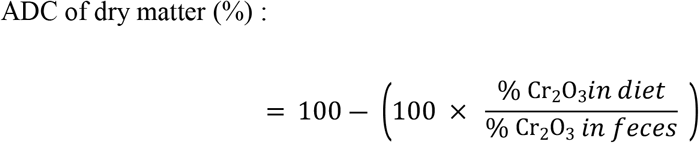

### 2.6. Chemical analysis

According to Horwitz and Latimer [10], samples of feed ingredients, feces, and diets were homogenized for analysis. Moisture was analyzed by drying the samples in oven at 105 °C for 12 hours. Kjeldhal apparatus was used for determination of crude protein (N*6.25) and Soxtec apparatus was used for measurement of crude fat by petroleum ether extraction method. While gross enegy was evluated by oxygen bomb calorimeter (Parr interment Co. Moline, USA.) and contents of chromic oxide were determined through calorimetrically at 720 and 350nm wavelength absorbance. According to methods devised by NRC [11], coefficient of apparent digestibility was determined.

### 2.7. Statistical analysis

Data obtained during present study was subjected to ANOVA using (SAS 9.1). The difference among means was compared by Duncan’s multiple range test.

## 3. Results

Table 3 and Table 4 lists the proximate analysis of the feces of juvenile L. rohita fed various experimental meals. The percentage of moisture in the fecal material of L. rohita fed fishmeal ranges from 21.25 to 26.89% in the juvenile given rusk cake. Fish fed biscuit waste had the highest crude protein content (60.44%) in their feces, whereas fish fed rusk waste had the lowest (50.75%). While there were non-significant variations in the level of lipids among the various regimens. On the other hand, fish fed diet containing fishmeal showed a maximum gross energy of 7.3 MJ/kg and a minimum gross energy of 4.66 MJ/kg when fed bread waste.

**Table 3.** Proximate analysis of experimental diets.

| Percentage | Z | A | B | C | D | E | F |
| --- | --- | --- | --- | --- | --- | --- | --- |
| composition (%) |  |  |  |  |  |  |  |
| Moisture | 8.32 | 10.1 | 14.4 | 9.15 | 7.16 | 14.22 | 8.08 |
| Crude protein | 51.48 | 41.76 | 40.89 | 39.18 | 42.15 | 41.94 | 41.22 |
| Crude lipid | 8.85 | 7.11 | 10.1 | 9.93 | 9.78 | 9.63 | 10.83 |
| Ash | 27.7 | 21.3 | 24.67 | 23.99 | 26.2 | 24.5 | 24.16 |
| Crude fiber | 0.68 | 0.99 | 8.3 | 4.68 | 5.43 | 3.57 | 5.79 |
| Chromic oxide | 1.34 | 0.41 | 0.63 | 1.54 | 0.56 | 1.52 | 1.35 |
| Gross energy (Mj/kg) | 15.64 | 14.32 | 17.68 | 16.53 | 17.68 | 15.73 | 16.27 |

\*Z=Control Diet-Fish Meal, A= Wheat Flour, B= Cake waste, C= Rice Polish, D= Biscuit waste, E= Bread waste, F=Rusk waste.

**Table 4.**
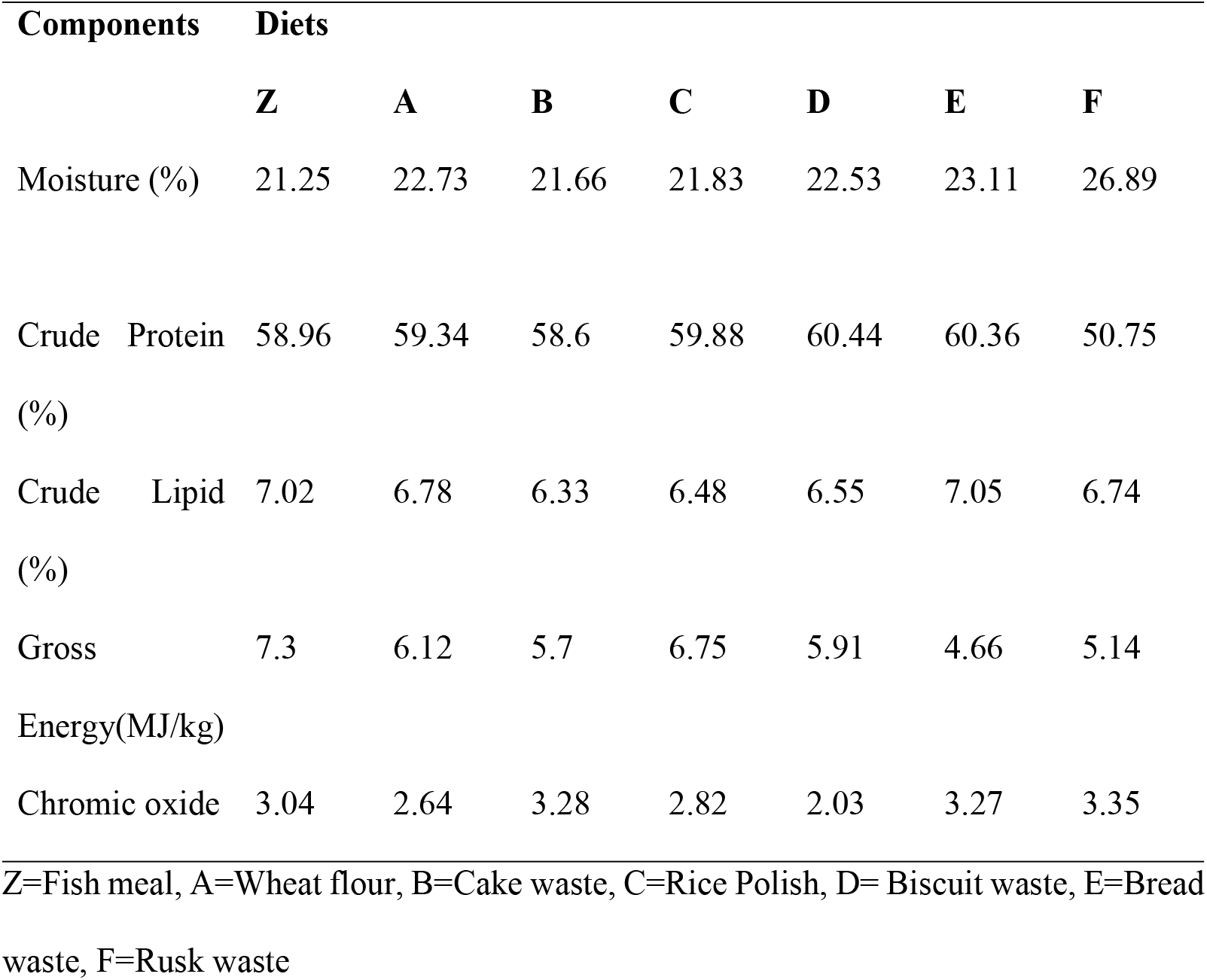
Proximate composition of feces of *L. rohita* juveniles fed with experimental diets.

| Components |  | Diets |  |  |  |  |  |  |
| --- | --- | --- | --- | --- | --- | --- | --- | --- |
|  |  | Z | A | B | C | D | E | F |
| Moisture (%) |  | 21.25 | 22.73 | 21.66 | 21.83 | 22.53 | 23.11 | 26.89 |
| Crude Protein (%) |  | 58.96 | 59.34 | 58.6 | 59.88 | 60.44 | 60.36 | 50.75 |
| Crude Lipid (%) |  | 7.02 | 6.78 | 6.33 | 6.48 | 6.55 | 7.05 | 6.74 |
| Gross Energy(MJ/kg) |  | 7.3 | 6.12 | 5.7 | 6.75 | 5.91 | 4.66 | 5.14 |
| Chromic oxide |  | 3.04 | 2.64 | 3.28 | 2.82 | 2.03 | 3.27 | 3.35 |

Z=Fish meal, A=Wheat flour, B=Cake waste, C=Rice Polish, D= Biscuit waste, E=Bread waste, F=Rusk waste

The growth performance of *L. rohita* fingerlings is displayed in Table 5. Group F given rusk cake had the highest mean weight gain (6.87 ± 0.30g), whereas group C fed rice polish had the lowest mean weight gain (4.53 ± 0.53g). However, with respect to average weight gain, notable variation observed among all treatments at p < 0.05. Group F had the highest percent weight increase value (156.88 ± 6.99%), while group C had the lowest value (111.26 ± 19.41%). For percentage weight gain, non-significant difference were determined across all treatments at p > 0.05. However, lowest (2.32 ± 0.04) values of FCR was noted in treatment F while highest (3.2 ± 0.15) was observed in fish fed with treatment C. Additionally, highly significant difference was determined for SGR percentage among all groups (p < 0.05).

**Table 5.**
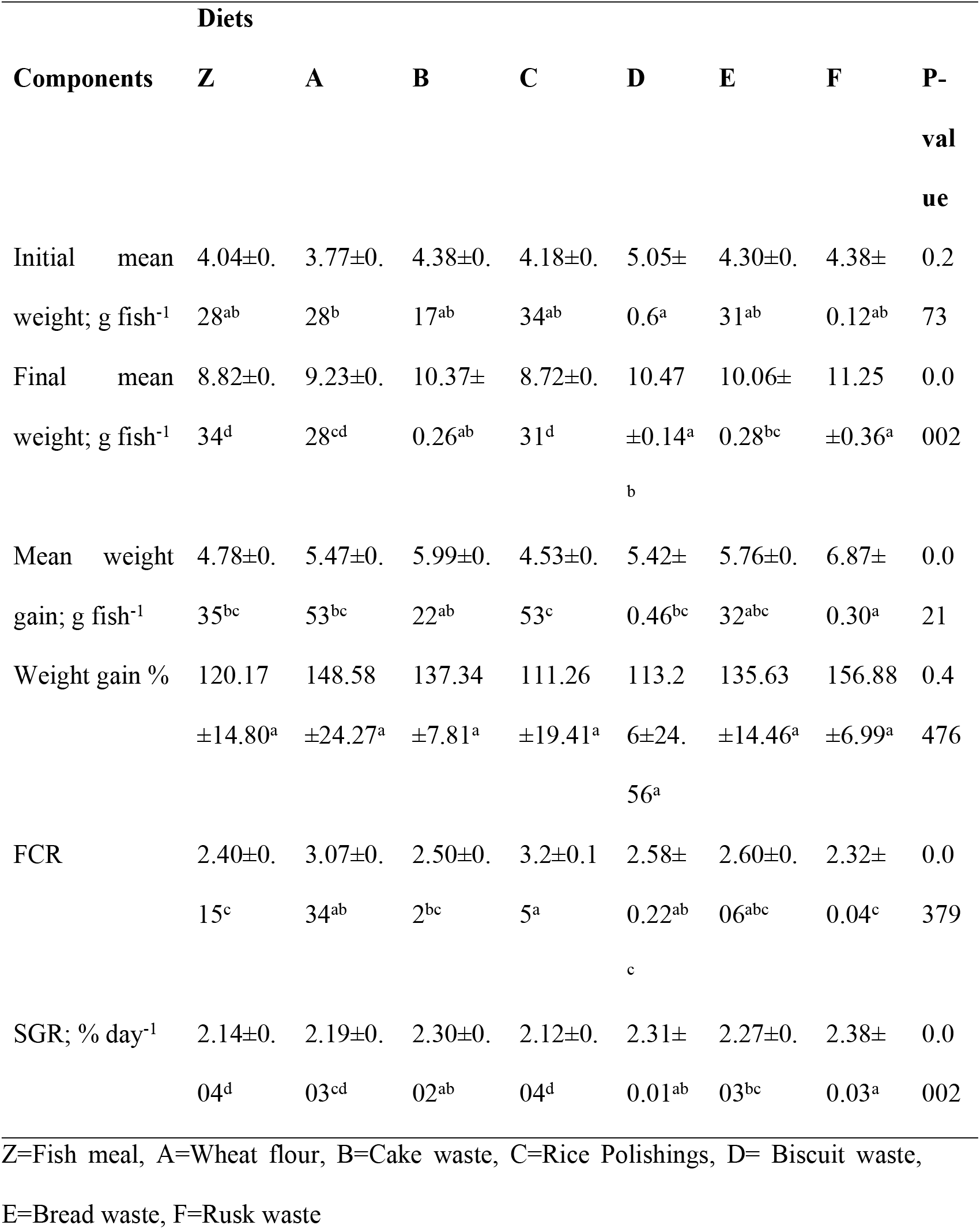
Growth parameters of L. rohita fed with diets of different test ingredients.

Figure displayed the ADC of gross energy, nutrients, and dry matter. The ADC values of protein differ significantly (p < 0.05) across all groups, with group A exhibiting the highest value (77.70 ± 0.05%) and group C displaying the lowest value (16.13 ± 1.86%). Similar findings regarding the protein’s ADC were displayed by groups A and B. Group B had the highest ADC value for the lipid (87.88 ± 0.14%), whereas groups A and C showed the minimum values (64.60 ± 0.27% and 64.21 ± 0.98%), respectively. For the average values of lipids, significant differences (p < 0.05) observed across all the groups. Groups A (8.29 ± 0.08) and Z (8.33 ± 0.05) had the lowest ADC of gross energy, while group B had the highest value (11.97 ± 0.02). Significant differences were seen in the overall results for the ADC of the gross energy (p < 0.05). Group F had the highest ADC of dry matter (70.14 ± 0.03%), whereas group D had the lowest (50.62 ± 0.05%). Overall, highly significant results were obtained for ADC among all treatments.

## 4. Discussion

For the formulation of nutrient rich diets, the evaluation of apparent digestibility coefficient is considered to be the most important factor [12,13]. When determining the effectiveness of feed ingredients, digestibility is the core factor which described the amount of nutrient available to body and not excreted as feces [8]. In accordance, important information required for effective formulation feed is digestibility of dry matter of nutrients [14].

The results of the current study revealed that significant differences were observed with respect to ADCs of protein, lipid, and dry matter. Variation among the digestibility of dry matter of different ingredients suggest that *L. rohita*, on the basis of kinds of ingredients, utilize them differently. In accordance to current findings, in terms of dry matter digestibility, gradual improvement with bakery waste determined in common carp fingerlings [15] and in lambs [16].

Carbohydrate digestibility, might be the reason for significantly enhance in digestibility of bakery waste in diets. Zhou and Yue [17] conducted a study on the hybrid tilapia and found significant difference for the average value of apparent digestibility of dry matter for ingredients based on plant and animal origin. They reported that hybrid tilapia respond to animal originated ingredients more effectively as compare to other. Inline to previous study, anti nutritional factors and high fiber contents in plant originated ingredients, might be the cause for the low digestibility [18,19].

The dry rusk, cream cake, biscuits, and bread that were discarded from bakeries showed a lower moisture content than the typical components found in feedstuff. Possible causes for this include the intense heat and finely crushed and sieved wheat used in bread-making processes, which facilitate quick drying. In the control diet and other bakery waste items such as biscuits, cream cake, bread, and dry rusk, the value of crude lipids was nearly same. The average values of crude protein was determined to be lowest in control group with respect to all other diets containing different bakery waste ingredients. Since fine, refined flour is used to produce bakery goods, the high energy ratio may be the cause of this difference. It is also challenging to compare prior research on the chemical composition of bakery waste, as reported by Franca et al., [20]. This is mostly due to variations in the quality and origin of the ingredients as well as in storage and processing techniques. The quality of an ingredient is determined by the availability of the quality and quantity on amino acids [21]. While, during present study, all formulated feed have all the essential amino acids including lysine, leucine, methionine, isoleucine, phenylalanine, histidine, tryptophan, arginine and valine, although in different proportions [22].

During present study, no adverse effect of use of bakery waste ingredients in the formulation of feed observed as it improved the growth performance of *L. rohita* fingerlings. Comparable outcomes were noted when maize meal and barley flour was used up to 75% in common carp fingerlings and substituted with bakery waste [23]. Therefore, bakery waste can be used as cheaper and healthier ingredient for the formulation of feed as it does not had any deleterious impact on the efficiency and growth rate of *L. rohita* fingerlings. Increase in average weight gain was observed in fingerlings of African giant catfish, when 30% of dietary maize was substitute with bread waste [24]. Similar results were observed in sail, when 22% of maize was replaced with bread waste followed by up to 100% replacement [25]. Additionally, these findings are in accordance with the studies conducted for terrestrial farmed animals. Hetherington and Krebs [26] conducted experiment on sheep to evaluate the effect of different dietary inclusion of bread waste such as 0, 25, and 50%. They found non-significant variations among different treatment for the average values of feed conversion ratio and live weight gain while the higest values were found in treatment fed with 50% inclusion.

Mahmoud [27] determine increase in growth rate with specific lamb diets, bakery by-products may be substituted with wheat bran and maize in concentrated feed mixes up to 60% of the ration. Al-Tulaihan at al., [28] examined how dried bread trash was incorporated into broiler diets with good success for the poultry industry. The earlier writers concluded that performance was unaffected by replacing around 50% of the dietary maize inclusion. Similarly, Japanese quail chicks’ meals might consist entirely of bakery waste instead of corn [29] and pig [30] without any harmful impact on growth rate.

The increase in digestibility of starch in bakery waste might be due to thermal processing (cooking) during the bread-making process may be the cause of the improvement in the digestion of carbohydrates for fish [27,30]. The starch granule is successfully broken down by gelatinization, which happens in the presence of heat and water. This increases the surface area of the starch exposes the bound amylase portion, making the starch more easily attacked by enzymatic action [31,32]. Gelatinization enhanced the digestibility of starch as compared to normal starches [33,34].

All these have attributes in diets for rohu due to its omnivorous mode of nutrition and capacity to digest and assimilate dietary starches and sugars as energy sources. It should be noted that the significantly lower protein digestibility observed for fish fed rice polish could be the consequence of protein-fibre interactions in the gastrointestinal tract for *L. rohita* affecting the ability of digestive enzyme action. This species has a reduced gastric function and relies mainly on the secretion of pancreatic proteases. Starch utilization in fish is complex and this is described by Salihu et al., [35] on their work to evaluate various starch rich flours in experimental diets for juvenile tinfoil barb (*Barbonymus schwanenfeldii*) Different types of starches with presence of fibre (bran) may also affect liver function and secretion of key digestive enzymes in the intestine.

More work is required to ascertain specific nutrient interactions of different processed carbohydrates within the composition of bakery waste products for warm water fish species.

Due to high digestibility and efficiency, bakery waste proved to be promising for formulation of feed for *L. rohita* fingerlings. The palatability of bakery wastes (cake waste, biscuit waste, bread waste, and rusk waste) along with their excellent nutritional value including appreciable levels of protein and lipids (fat and oil) offers much potential. Present finding provides useful insights for the formulation of nutritionally balanced as well cost effective feed with the help of bakery waste such as cake waste, biscuit waste, bread waste, and rusk waste. In the context of the developing aquaculture sector in Pakistan involving several species of economic importance, this preliminary investigation with rohu provided useful information for feed formulation using such byproducts. This research needs to be applied to various phases of *L. rohita* production beyond juveniles and expanded to other species such as rainbow trout, pangasius, tilapia and even shrimp. In this manner, importation of major ingredients may be reduced and offering a more sustainable and resilient aquafeed sector for Pakistan.

---

**Figure.**
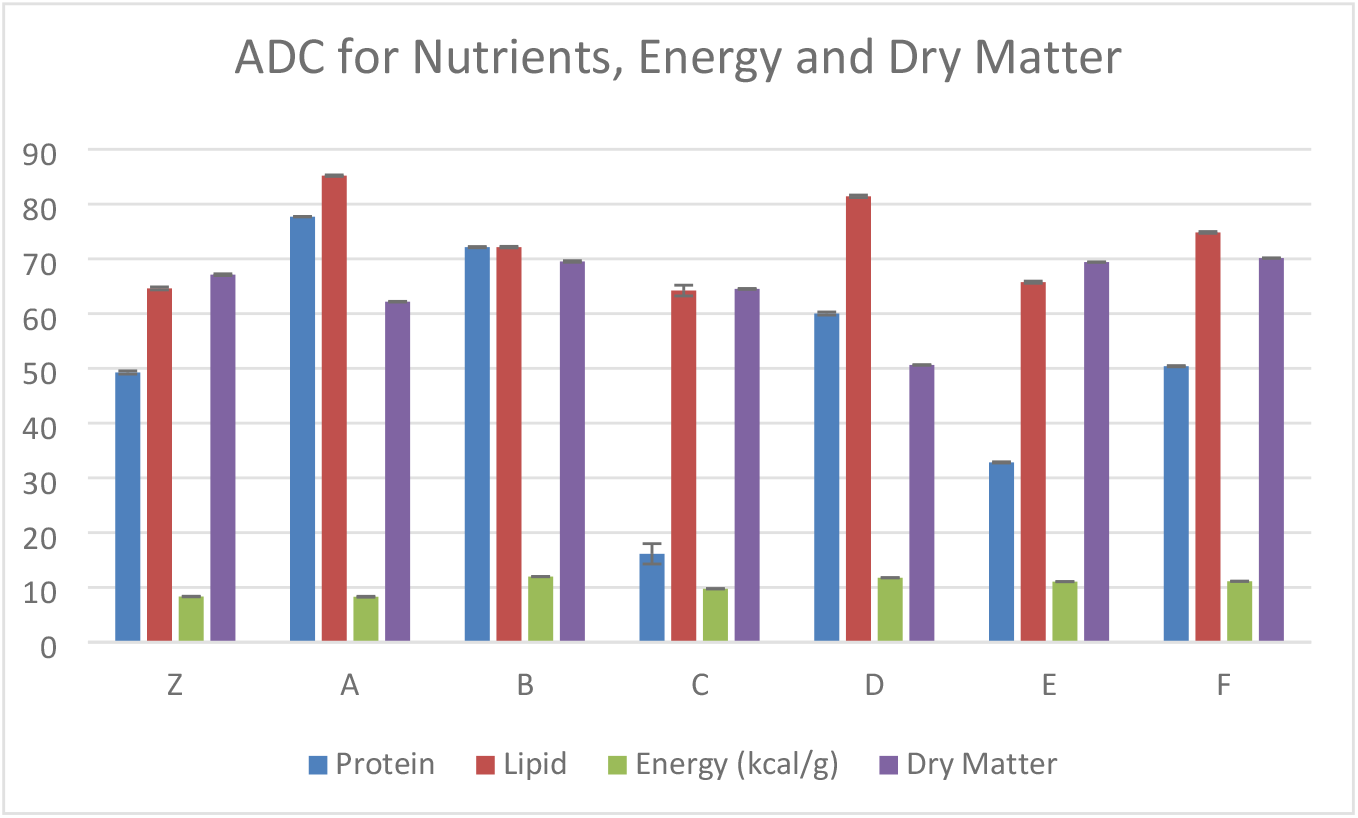
ADC for nutrients-protein, lipid, energy and dry matter

